# Mechanical Activation of Piezo1 by Virus-like Nanospikes to Potentiate STING-driven Macrophage Reprogramming

**DOI:** 10.64898/2026.08.05.742741

**Authors:** Jiajia Wang, Minna Sivonen, Enkhzaya Batnasan, Sini Pitkänen, Janne Tampio, Adela Králová, Minna-Mari Tervo, Leena Latonen, Anna-Liisa Levonen, Kristiina M. Huttunen, Tarja Malm, Rashid Giniatullin, Vesa-Pekka Lehto, Wujun Xu

## Abstract

Mechanotransduction plays a fundamental role in regulating immune cell function, yet how engineered virus-like nanospikes engage mechanosensitive signaling pathways to modulate innate immunity remains poorly understood. Here, we report virus-like nanotopography as a previously unrecognized regulator of Piezo1-mediated mechanotransduction in macrophages using virus-like mesoporous silica nanoparticles (VLPSi) with tunable rigid nanospike lengths. We demonstrate a direct structure–activity relationship between nanospike geometry and Piezo1-dependent Ca²^+^ influx, with longer nanospikes inducing significantly greater intracellular Ca²^+^ signaling. Building on this mechanistic insight, we developed biomimetic cancer cell membrane (CM)-coated, MSA-2-loaded VLPSi nanoparticle (CM/MSA-2@VLPSi) and investigate the combination of nanospikes-activated Piezo1 with STING signaling and CM antigens presentation in macrophage immune reprogramming. The resulting biomimetic nanoparticles robustly activate the STING–TBK1–IRF3/NF-κB axis, increase IFN-β and pro-inflammatory cytokine production, and promote macrophage polarization toward M1 phenotype in a spike-length-dependent manner. Collectively, the present study provides a biomimetic strategy for enhancing the M1 polarization of macrophage through the coordinated regulation of mechanical, inflammatory, and antigenic signals.

## Introduction

Macrophages are multifunctional cells of the innate immune system that participate in both inflammatory responses and the maintenance of tissue homeostasis^1^. Resting macrophages (M0) exhibit notable plasticity and can polarize into special phenotypes in response to environmental stimulation, primarily the classically activated M1 phenotype and the alternatively activated M2 phenotype^2^. M1 polarized macrophages are considered pro-inflammatory cells involved in host defense. The M1 phenotype is characterized by activation of NF-κB and interferon regulatory factor 3 (IRF3), leading to the production of pro-inflammatory cytokines such as TNF-α, IL-6, and IL-1β, together with high expression of CD80. In the tumor microenvironment, M1 macrophages are regarded as protective and antitumorigenic^3^. Moreover, M1 macrophages promote Th1 immune responses, thereby having a positive effect that enhances antitumor immunity^4^.In contrast, Th2 immune responses are typically associated with activated M2 macrophages. M2 macrophages are anti-inflammatory, not only involved in tissue repair and wound healing but also boost tumor progression. M2 polarization is induced by IL-4 and increased IL-10 secretion, and high expression of CD206^5^. Therefore, M1 macrophages promote antitumor immunity through enhanced inflammatory signaling, antigen presentation, and activation of adaptive immune responses, making M1 polarization a desirable therapeutic goal.

Mechanotransduction plays a fundamental role in regulating immune cell functions, such as macrophage programing^6^. Piezo 1 is a mechanosensitive, non-selective cation channel located on the plasma membrane. It plays a critical role in sensing and transducing external mechanical stimuli into electrochemical signals that regulate intracellular signaling pathways and cellular behavior^7^. Activation of Piezo1 typically induces intracellular Ca²^+^ influx, a fundamental signaling event that regulates diverse cellular processes. Disruption of Ca²^+^ homeostasis can trigger different cellular stress responses and serves as a key initiator of immune signaling cascades^8^. Previous studies have investigated Piezo1 activation through magnetomechanical stimulation^9^, particle stiffness engineering^6,10^, and chemical redox regulation^11^. Despite growing interest in Piezo1-mediated mechanotransduction, the ability of bioinspired nanospike architectures to activate Piezo1 signaling and modulate downstream immune responses remains largely unexplored.

In recent years, the stimulator of interferon genes (STING) has emerged as another attractive target for activating immunity in cancer therapy^12,13^. MSA-2, a small-molecule STING agonist, activates the cyclic GMP–AMP synthase (cGAS)–STING pathway and leads to initiate IFN-I expression. This signaling cascade induces the production of type I interferons, pro-inflammatory cytokines, and chemokines and then enhances antigen cross-presentation by mature antigen-presenting cells (APCs) for activating immune response^14^. In addition to its immunostimulatory effects, activation of the STING pathway has been shown to reprogram immunosuppressive M2-like tumor-associated macrophages (TAMs) toward a pro-inflammatory M1 phenotype, characterized by increased CD86 expression, enhanced inflammatory signaling, and improved antitumor immune response^15^. Given the critical role of M1 macrophages in promoting Th1 immunity and suppressing tumor progression, STING agonists have emerged as promising candidates for macrophage-targeted cancer immunotherapy. However, the promising adjuvant effects of STING agonists in cancer vaccines, systemic administration may result in “on-target, off-tumor” STING activation, rapid clearance, low bioavailability, and poor specificity^16^. Therefore, the development of efficient delivery platforms capable of enhancing STING agonist accumulation within target immune cells remains highly desirable. Most of the synthetic nanoparticles (NPs) are easily recognized by macrophages, making macrophages act as key regulators in nanomaterial-mediated immune response. Importantly, the physicochemical properties of nanomaterials, including size^17^, surface roughness^18^,and surface functionalization^19^, significantly influence macrophage uptake efficiency and polarization status^20,21^. Therefore, NPs have been widely studied as promising nanocarriers in biomedical applications^22^. In particular, virus-like mesoporous silica (VLPSi) NPs composed of an internal spherical mesoporous silica core surrounded by vertically arranged spiky silica nanotubes have been synthesized and applied as nanoplatforms for drug delivery^23^. Our recent study longer spikes nanoparticles significantly enhanced cellular uptake and promoted the maturation of monocyte-derived dendritic cells (MoDCs) and upregulated key maturation and antigen-presentation markers, including CD86, CD40, and HLA-DR, indicating spike-length– dependent immune activation. Mature DCs further stimulated stronger T-cell proliferation and division^24^. These results demonstrate that rigid spike topology enhances DC activation and subsequent T-cell responses, highlighting the potential of spike-engineered VLPSi as immunomodulatory platforms for improved immunotherapy^25^.

The aim of this study is to elucidate how virus-like nanotopography regulates macrophage immune responses through Piezo1-mediated mechanotransduction and whether this process can be leveraged to enhance STING-driven immunotherapy (Scheme 1). To this end, we developed a biomimetic platform by loading the STING agonist MSA-2 into VLPSi and subsequently coating them with cancer cell membranes (CMs) derived from IFN-γ-treated tumor cells. To investigate the role of nanotopography in Piezo1 regulation, VLPSi with distinct spike lengths (VLPSi-5 and VLPSi-30) were synthesized and systematically evaluated. Furthermore, we examined whether the integration of virus-like nanotopography, STING activation, and biomimetic CM coating could synergistically modulate innate immune responses. The biomimetic platform succeeded in efficiently boosting M1 macrophage polarization via the combined effect of enhanced cellular uptake, evoked Ca^2+^ influx, activated STING pathway, and improved antigen presentation. Collectively, the present study provides a bioinspired framework for integrating mechanical, inflammatory, and antigenic signals to boost macrophage-based immunotherapies.

**Scheme 1.**
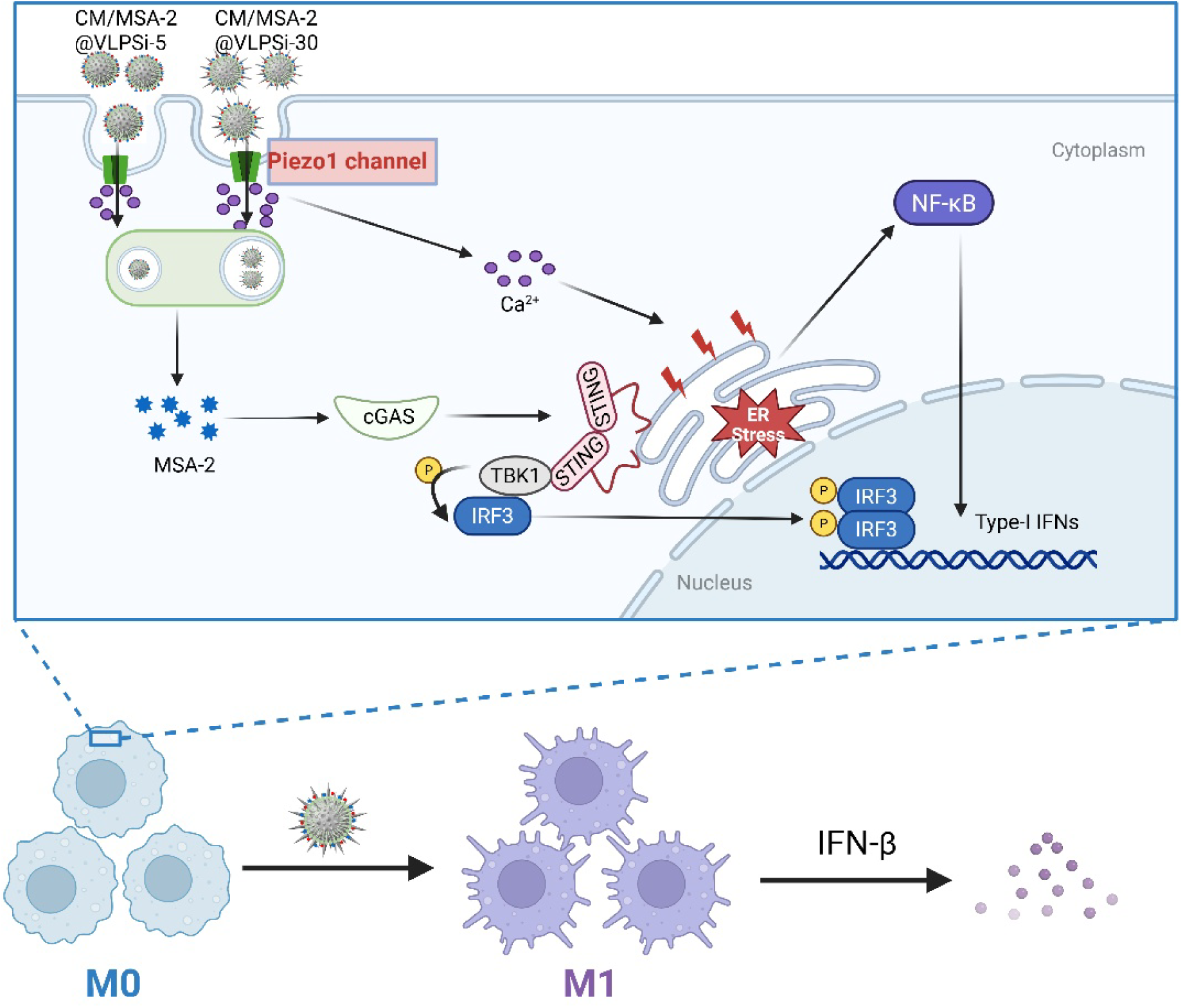
Schematic illustration of mechanical activation of Piezo1 to potentiate STING-driven macrophage reprogramming. Rigid nanospikes on VLPSi enhance Piezo1-dependent Ca²^+^ influx and activate NF-κB signaling, while intracellularly released MSA-2 activates the STING pathway promote TBK1/IRF3 phosphorylation, type I interferon production, and macrophage polarization.

## 1. Materials and methods

### 1.1 Preparation of the virus-like silica nanoparticles

1% Hexadecyltrimethylammonium bromide (CTAB, Sigma) was mixed with 0.1 M sodium hydroxide in deionized water at 60 °C. A mixture of 20 mL tetraethyl orthosilicate (TEOS, Merck) and 80 mL cyclohexane (Merck) was then added by Aladdin Single-Syringe infusion Pump (World Precision Instruments). The spike length of the VLPSi was controlled by adjusting the reaction time: 40 h for 5 nm spikes and 70 h for 30 nm spikes. Following the reaction, the VLPSi NPs were collected by centrifugation, dried in the oven, and then calcined in the Tube Furnaces (Nabertherm GmbH) at 600°C in air atmosphere for 4h.

### 1.2 Cell lines

RAW 264.7 cell lines were cultured in Dulbecco’s modified Eagle’s medium (DMEM, VWR) medium supplemented with 10% fetal bovine serum (FBS, Gibco) and 1% Penicillin-Streptomycin-Glutamine (Gibco). B16-F10-OVA cells were cultured in 10% fetal bovine serum supplemented RPMI 1640 (VWR) and 0.4 mg/mL Geneticin (G418, InvivoGen). Trypsin-EDTA (0.25%), phenol red (Gibco)and TrypLE™ Express Enzyme (1X), phenol red (Gibco) were used to detach the cells. Cells were maintained at 37 °C in a humidified atmosphere containing 5% CO_2_.

The human immortalized microglial cell line SV-40 (T0251, Applied Biological Materials Inc.), and kindly provided by Dora Brites (University of Lisbon, Portugal)^26^, was cultured in DMEM□+□GlutaMAX (4.5 g/L glucose; Gibco) with 10% FBS and 1% penicillin/streptomycin. Cells were grown and seeded at 40,000 cells/cm² to reach approximately 80% confluence after 24 hours for experiments.

Human induced microglia-like cells (iMGLs) were differentiated from the wild type (WT) iPSC line GM25256 and the Piezo1 knockout clone F6 generated from the same parental line at Tampere University. Briefly, iPSCs were dissociated into single cells and cultured on Matrigel-coated plates in E8 medium supplemented with BMP4, Activin A, CHIR99021, and ROCK inhibitor Y-27632 under hypoxic conditions. Cells were subsequently differentiated into erythromyeloid progenitors (EMPs) using differentiation medium containing FGF2, VEGF, IL-3, SCF, IL-6, and thrombopoietin. Floating EMPs were collected at day 8 and transferred into ultra-low attachment dishes in microglial differentiation medium supplemented with insulin, M-CSF and IL-34. On day 10, progenitor cells were cryopreserved in Bambanker freezing medium (BB05, VingLab) and stored in liquid nitrogen. For experiments, frozen progenitors were thawed in medium containing ROCK inhibitor Y-27632 and further matured in microglial maturation medium with M-CSF and IL-34. Cells were plated in 35 mm cell culture dishes (Sarstedt) at day 15 and used for experiments at day 21 and 25. Detailed differentiation procedures were performed as previously described^27^.

### 1.3 Collection of cell membrane

To enhance MHC-I expression levels of B16-F10-OVA cell surface, 1.5×10^4^ per well B16-F10-OVA cells were stimulated with 0, 25, 50, 100 ng/mL IFN-gamma (IFN-γ, Fisher Scientific Oy) for 24 h. After washing with PBS (VWR), the cells were stained (15 min +4°C) with zombie green^TM^ fixable viability kit (Biolegend) according to manufacturer’s instructions. Cells were subsequently stained in flow buffer (0.02 % NaN3, 2 mM EDTA, 2 % FBS in PBS) with anti-mouse H-2 Antibody-PE (Biolegend). Cells were measured with CytoFLEX S (Beckman Coulter) and analyzed with CytExpert v2 (Beckman Coulter) and FlowJo v10 (BD Life Sciences, TreeStar). After the MHC-I expression was confirmed, the resultant cells were then collected to isolate cell membranes using a previously described method^28^. Briefly, IFN-γ stimulated cells were suspended in a hypotonic lysis buffer (PH=7.5) containing 20 mM Tris-HCl (Sigma-Aldrich), 10 mM KCl, (Merck), 2 mM MgCl2 (Merck) and EDTA-free protease inhibitor (one tablet for 10 mL buffer solution, Roche Diagnostics GmbH), and incubated on ice for 30 min. Dounce pressure homogenization method was used. The obtained supernatants were centrifuged at 100000 rcf at 4 °C. Then, the cell membrane protein concentration was quantified by Pierce™ Dilution-Free™ Rapid Gold BCA Protein Assay Kit (Thermo Scientific) using VICTOR Nivo Multimode Microplate Reader and stored in 1 mL of HEPES buffer (25 mM HEPES, pH = 7) at -20 °C.

### 1.4 Preparation of CM/MSA-2@ VLPSi

VLPSi NPs were mixed with 2-[methoxy(polyethyleneoxy)propyl] trimethoxysilane (90%, 6–9 PEG units, abcr) in ethanol and heated to 90 °C in an oil bath^29^. After 30 mins of stirring, the VLPSi NPs were successfully PEGylation and got samples PEG-VLPSi-5, PEG-VLPSi-30.

VLPSi NPs (including VLPSi-5 and VLPSi-30) and N-[3-(Trimethoxysilyl) propyl]-N, N, N-trimethylammonium chloride (50% in methanol, thermos scientific) were mixed with volume ratio=5:1. After washing twice, the sample PEG-VLPSi-N(CH_3_)_3_-5 and PEG-VLPSi-N(CH_3_)_3_-30 were prepared. MSA-2 (MedChemExpress) was sonicated with PEG-VLPSi-N(CH_3_)_3_ particles for 30 mins with volume ratio=1:5 and then got MSA-2@ VLPSi-5, MSA-2@ VLPSi-30. The MSA-2 loading efficiency was determined using the equation.

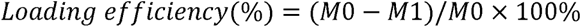

where M0 is the initial amount of MSA-2 before loading; M1 is the amount of free MSA-2 detected in the supernatant.

MSA-2@ VLPSi particles were mixed with CMs at weight ratio of membrane protein/NPs =1/5. Then, the mixtures were sonicated in 2 mins and mixed in solution (PBS: DDH_2_O = 1:1) at room temperature for 40min by rotator. After washing, the fresh sample CM/MSA-2@ VLPSi-5 and CM/MSA-2@ VLPSi-30 was got and used immediately.

### 1.5 Physicochemical characterizations

Diluted suspensions of VLPSi were imaged on the Carbon Film 400 Mesh Cu (HC400-Cu, USA). The particle morphology was characterized by transmission electron microscopy (Jeol JEM-2100F TEM). The zeta potential of samples was measured by using Zetasizer Nano ZS (Malvern Panalytical Ltd, United Kingdom).

Thermogravimetric analysis (TGA) analyses were performed using a Netzsch TG 209 F1 Libra instrument, The samples were analyzed under N_2_ gas flow, from 100□°C to 900□°C using a heating rate of 20□°C/min.

### 1.6 Detection of intracellular calcium ion content

Ca^2+^-imaging was performed in SV40-immortalized microglial cells, induced microglia (iMGs) derived from the wild-type (WT) iPSC line GM25256 and the Piezo1 knockout clone F6 following previous procedure^30^. Cells were divided into cell culture dishes (35 mm diameter, Sarstedt) and then incubated in fresh basic salt solution (BSS,152 mM NaCl, 10 mM HEPES, 10 mM glucose, 5 mM KCl, 4 mM CaCl_2_, 2 mM MgCl_2_, pH = 7.4) containing 2 µm Fluo-4 Direct™ Calcium Assay Kit (Invitrogen) at 37°C for 30 min. It was followed by washing twice with BSS before adding the samples of VLPSi-5, VLPSi-30 (100µg/mL) with BSS in the cell dishes. After that the Ca^2+^-imaging was performed at one frame per second for 30 mins using 10x objective (Olympus IX-7010). Ionomycin calcium salt (Tocris) was applied in the 1700 second as positive control. Fluorescence was detected using the Till Photonics imaging system (FEI GmbH) equipped with a 12-bit CCD Camera (SensiCam) with an excitation wavelength of 495 nm. Regions of interest (ROIs) around the cell body were selected from the whole image with Image J software. The average of the first 5 minutes of the recording was used as a baseline. Fluorescence intensity change over time is defined as ΔF/Fo = (F − Fo)/Fo, where F is the fluorescence intensity at any time point, and Fo is the baseline fluorescence intensity averaged across 5 minutes from the beginning for each cell. Structural visualization of mouse Piezo1 based on the high-resolution cryo-electron microscopy (cryo-EM) structure 6B3R by PyMOL^31^. Structural models from cryo-EM were obtained from the Protein Data Bank (PDB)^32^.

### 1.7 Release profile of MSA-2

*In vitro* release of MSA-2 from CM/MSA-2@VLPSi was conducted. The samples were dispersed and rotated into 1 ml PBS buffer solutions at 37 °C. At different time points (0, 2, 4, 6, 8, 24h), 0.5 ml of the supernatant was collected by centrifugation, and the pellet was resuspended with 0.5 ml fresh PBS to maintain the system at 1 ml. The obtained supernatant was diluted with 50% ACN 1:20 (v/v) and analyzed with an ultra-high performance liquid chromatography (UHPLC; Agilent Technologies 1290 Infinity II system; Agilent Technologies Inc., Wilmington, DE, USA). The instrument consisted of a high-speed pump (G7120A), a multisampler (G7167B), a multi-column thermostat (G7116B), and a diode array detector (G7117B). The chromatographic separation was performed using a ZORBAX Eclipse Plus C18 column (2.1 x 50 mm, 1.8 µm; Agilent Technologies Inc., Wilmington, DE, USA) and eluents of A) H2O with 0.1% formic acid and B) ACN with 0.1% formic acid in an isocratic ratio of 70:30 (A: B). The mobile phase flow speed was 0.3 ml/min, the column compartment was set to 25 °C, and the injected sample volume was 2 µl. The detection wavelength for MSA-2 was 320 nm. The MSA-2 concentrations from the samples were calculated using a standard curve (0.5-10 µg/ml).

### 1.8 Cell cytotoxicity and cellular uptake

The cytotoxicity of CM/MSA-2@ VLPSi-5 and CM/MSA-2@ VLPSi-30 was detected using ATP assay kit. RAW 264.7 cells (1.5 × 10^5^ cells per well) were incubated in a CO_2_ incubator containing 5% CO_2_ at 37 °C overnight. 25, 50, 100, 200 µg/mL CM/MSA-2@ VLPSi-5 and CM/MSA-2@ VLPSi-30 were added to each well and then incubated for 24□h. 50 μL fresh cell culture medium and 50 µL Cell Titer-Glo™ Luminescent Cell Viability Assay Kit (Promega) were added to each well and incubated at room temperature for 10min, and the values were recorded on Fluoroskan Ascent FL microplate reader (Thermo Labsystems).

To study cellular uptake, RAW 264.7 cells were incubated overnight in Ibidi 8-well plate at a density of 2 × 10^4^ cells per well. The materials CM/MSA-2@ VLPSi-5 and CM/MSA-2@ VLPSi-30 were labeled with Fluorescein 5(6)-isothiocyanate (FITC, Alfa Chemical) and then were incubated with cells for 4□h (100 μg/ml). The cells were then washed 3 times with PBS buffer. Cell Mask™ red plasma membrane staining solution at 5µg/mL was added and incubated with cells at 37□°C for 10□min. The cells were washed and fixed with 4% paraformaldehyde (Sigma-Aldrich) for 10 min at room temperature. 10 µg/mL DAPI (4′,6-diamidino-2-phenylindole, Sigma-Aldrich) were used to stain nucleus for 10 min at room temperature. Fluorescence images were photographed by confocal laser scanning microscope instrument (Zeiss LSM 700).

### 1.9 IFN-**β** Detection with ELISA

RAW 264.7 macrophages were seeded on a 12-well plate and grown at a cell density of 3 × 10^5^ cells per well and treated with 100 µg/mL of CM/MSA-2@ VLPSi-5 and CM/MSA-2@ VLPSi-30, non-treated cells as control. After 24 h, the cells were centrifuged at 300 x g for 5 min to keep the supernatant for quantification of IFN-β by using LumiKine^TM^ Xpress mIFN-β 2,0 ELISA kit (InvivoGen), according to the manufacturers’ protocols. Briefly, standards and samples (cultured medium) were added to each well of the mIFN-β capture antibody-precoated plates and incubated. After washing, it was incubated with Lucia-conjugated detection antibodies. The plates were washed again and incubated with QUANTI-luc^TM^ 4 reagent solution under light-protected conditions and then luminescence was measured by using a VICTOR Nivo Multimode Microplate Reader (Revvity).

### 1.10 SDS-PAGE and western blot analysis

The cell membrane proteins of CM/MSA-2@ VLPSi-5, CM/MSA-2@ VLPSi-30, MSA-2@ VLPSi-5, MSA-2@ VLPSi-30 and pure CM were characterized by the sodium dodecyl sulfate–polyacrylamide gel electrophoresis method (SDS-PAGE). After protein concentration was measured by Tecan Infinite M200, the SDS-PAGE sample loading buffer (4× Laemmli sample buffer) was added to the samples and the mixture was heated to 95 °C. Based on the instruction of manufacturer, samples were added and imaged into a 10% SDS-PAGE gel by using a Nitrocellulose-membranes (Thermo Fisher Scientific).

The RAW264.7 cells were seeded into a tissue culture dish 6 cm in 4 × 10^6^ cells/dish for 24 h. The cells were treated with 100 µg/mL CM/MSA-2@ VLPSi-5 and CM/MSA-2@ VLPSi-30 for 12 h. Untreated group was used as control. RAW264.7 cells were collected and lysed by Triton-X lysis buffer. Protein lysates were resolved by SDS-PAGE and electro transferred to a nitrocellulose membrane. The primary antibodies were STING (D1V5L) Rabbit mAb (CST-50494T, Cell Signaling Technology, 1:1000), TBK1/NAK (D1B4) Rabbit mAb (CST-3504T, Cell Signaling Technology, 1:1000), anti-NF-kB (p65), pAb (ADP-AG-25T-0004-R100, AdipoGen, 1:1000), IRF-3 (D83B9) Rabbit mAb (CST-4302T, Cell Signaling Technology, 1:1000) and Lamin polyclonal rabbit 1:5000 (Lamin B1, Abcam, Ab16048, Cambridge, UK). The anti-mouse or anti-rabbit HRP-conjugated secondary antibodies produced in goats (Invitrogen, Thermo Fisher Scientific) were used. Chemiluminescence reactions were generated using Clarity Western ECL Substrate reagent (Bio-Rad Laboratories) and ChemiDoc MP Imaging system (Bio-Rad Laboratories) was used to detect the signal.

### 1.11 Reverse transcription quantitative PCR (RT-qPCR)

To evaluate the inflammatory gene expression, RT-qPCR was performed. RAW 264.7 macrophages were seeded in a 12-well plate and allowed to grow at a seeding density of 4 × 10^5^ cells per well before being treated for 24 h. And then each group was treated with cell medium or 100 µg/mL of CM/MSA-2@ VLPSi-5 and CM/MSA-2@ VLPSi-30 respectively for 6 h followed by rinsing with PBS three times. RNA was extracted using High Pure RNA Isolation Kit (Roche 11828665001) and cDNA was synthesized using Transcriptor First Strand cDNA Synthesis Kit (Roche 04897030001) by using the manufacturer’s protocols. Then, real-time PCR was performed and quantified on PrimeTime Std qPCR Assays (Integrated DNA Technologies) and Universal Probe Library (Roche) on QuantStudio 3 Real-Time PCR system (Applied Biosystems). The sequences of primers for IL-1β, IL-6, TNF-a, Arg1, IL-4, and TGF-β were listed as follows Table 1. The relative expression of each gene was analyzed, and the levels of target gene mRNA were normalized to those of Ppia using the 2^−ΔΔ*C*^_T_.

**Table 1.**
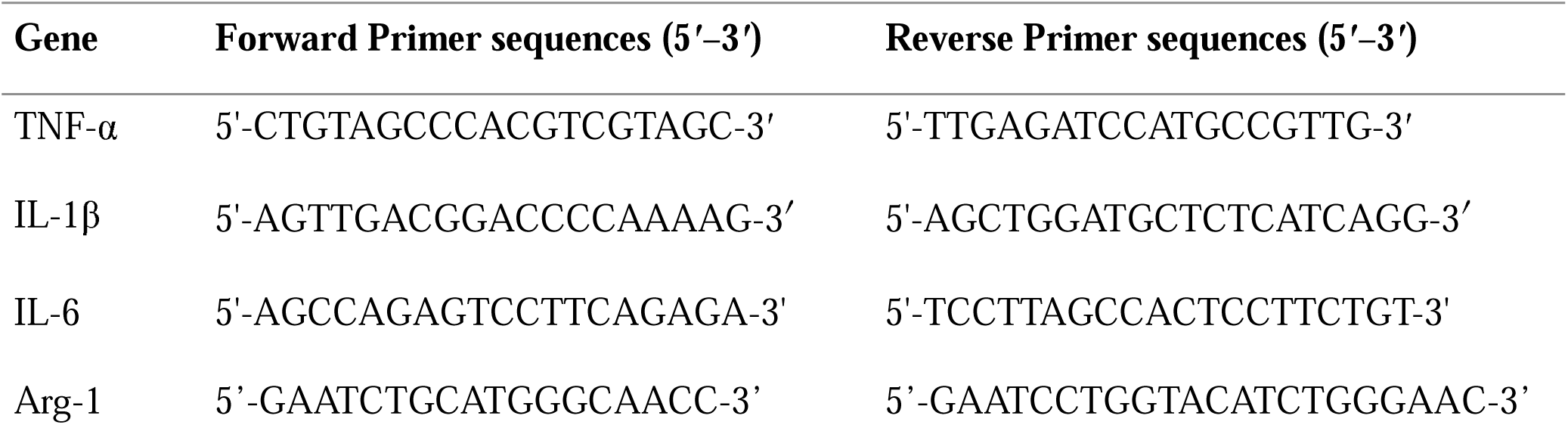
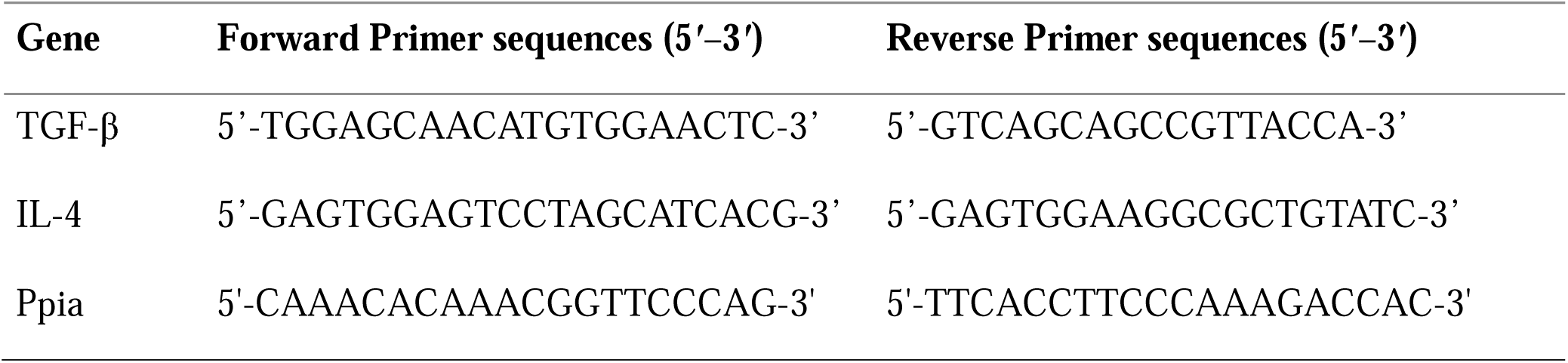
Primer sequences.

### 1.12 Flow cytometry to examine M1 and M2 polarization

RAW 264.7 cells were seeded in 12-well plates at 3×10^5^ cells per well and incubated overnight. Then, cells were stimulated with 100 ng/mL LPS + 20 ng/mL IFN-γ, 100 µg/mL CM/MSA-2@ VLPSi-5, CM/MSA-2@ VLPSi-30 for 24 h. Cells were rinsed with PBS (VWR) twice and incubated with zombie green^TM^ fixable viability kit for 15 min at 4 °C. After staining, the cells were washed and centrifuged at 300 x g for 5 mins. Cells were stained in flow cytometry buffer (0.02% NaN3, 2mM EDTA, 2% FBS in PBS) with PE-conjugated anti-mouse CD86 (M1 marker), APC-conjugated anti-mouse CD206 (M2 marker) antibodies for 15 min at room temperature. After incubation, the samples were washed twice with flow cytometry buffer and resuspended in buffer for analysis. Cells were acquired on CytoFLEX S (Beckman Coulter) and analyzed with CytExpert v2 (Beckman Coulter) and FlowJo v10 (BD Life Sciences, TreeStar).

### Statistics and reproducibility

Data were expressed as mean ± SD. All statistical analyses were performed using GraphPad Prism 10.4.2 software (GraphPad Software). A one-way ANOVA analysis was used to determine the statistical significance. The p-value is set in figures and legends such as: *P□<□0.05, **P < 0.01 and ***P□<□0.001.

## 2. Results and discussion

### 2.1 Preparation of VLPSi and physicochemical characterization

To deliver MSA-2 and CMs to cells, we developed VLPSi as a platform for nanocarrier. VLPSi was synthesized by using CTAB as a template to create a porous structure and TEOS as the silica precursor. TEM image confirmed the successful synthesis of VLPSi-5 and VLPSi-30, which exhibited uniform spike length, with an average spike length of approximately 5 nm and 30nm (Figure 1a-b). Subsequently, PEG was chemically conjugated on VLPSi to enhance biocompatibility of the NPs. TGA analysis showed that the weight of the VLPSi-5 and VLPSi-30, PEG-VLPSi-5 and PEG-VLPSi-30 decreased to 96.94%, 96.64%, 92.60% and 91.92% respectively, when the temperature reached 900 °C (Figure 1c). This indicated that approximately 4.34% and 4.72% of the PEG amount were grafted onto the VLPSi-5 and VLPSi-30.

**Figure 1.**
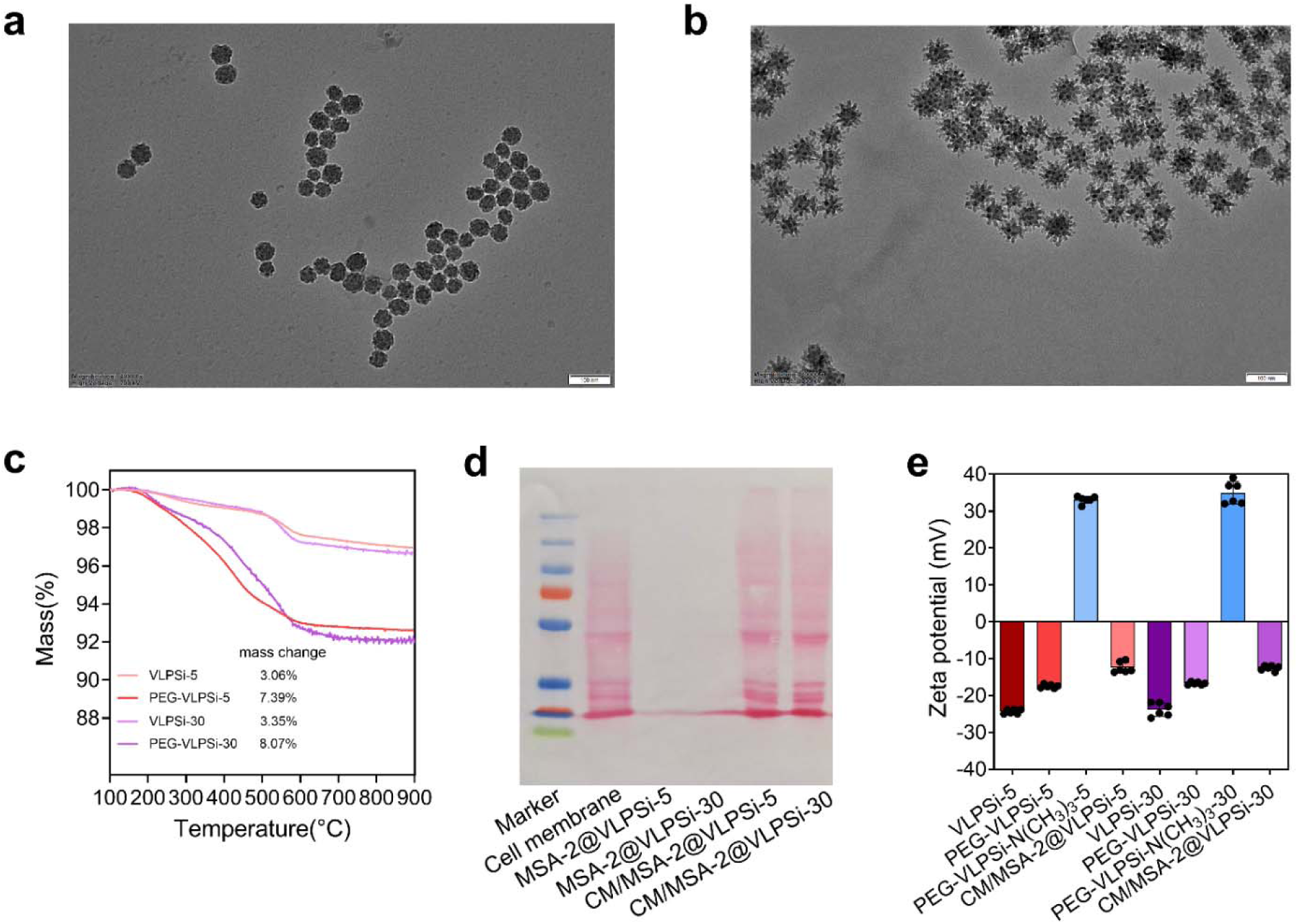
Physicochemical characterizations of VLPSi. a, b. Typical TEM images of the synthesized VLPSi-5 and VLPSi-30. Scale bars, 100 nm. c. Thermogravimetric analysis (TGA) curves of VLPSi to quantify PEG mass. d. SDS-PAGE analysis of cell membrane proteins. e. Zeta potential of VLPSi, PEG-VLPSi, PEG-VLPSi-N(CH_3_)_3_, CM/MSA-2@VLPSi

To study the effect of IFN-γ stimulating on the expression of MHC I, cells were treated with three concentrations (0, 25, 50, and 100 ng/mL). The best concentration of IFN-γ in promoting MHC I expression was observed at 25 ng/mL, and higher concentrations did not further change MHC I expression (Figure S1a, Supporting information). Hence, 25 ng/mL was used for subsequent experiments. Beyond serving as a marker of immune activation, the elevated MHC-I expression may facilitate the retention of tumor-associated peptide–MHC complexes and other immunoregulatory membrane components. These features could enhance the recognition and uptake of the NPs by macrophages, promoting their polarization toward the pro-inflammatory M1 phenotype. Importantly, activated M1 macrophages possess enhanced antigen-processing and antigen-presenting capabilities, which may further stimulate adaptive immune responses through the activation of tumor-specific T cells. Therefore, the IFN-γ-treated CM coating may contribute not only to innate immune activation but also to the initiation of a broader antitumor immune response. The loading efficacy of CM was investigated using SDS-PAGE and the BCA method. Based on gel retardation assays, the CM/MSA-2@VLPSi-5 and CM/MSA-2@VLPSi-30 showed successful CM loading (Figure 1d). The average CM loading efficacy of CM/MSA-2@VLPSi was 91.59% and 91.67%, respectively. The zeta potential of bare VLPSi-5 and VLPSi-30 in water was −24.35 mV and −23.83 mV and then changed to −17.48 mV and −16.72 mV respectively after modified with PEG (Figure 1e). Next, amine group -N(CH_3_)_3_ was modified on the PEG-VLPSi to load MSA-2 via electrostatic interaction. Zeta potential changed to +33.12 mV and +34.95 mV for PEG-VLPSi-N(CH_3_)_3_-5 and PEG-VLPSi-N(CH_3_)_3_-30. After CM and MSA-2 were loaded, the zeta potential of PEG-VLPSi-N(CH_3_)_3_ changed to -12.43 mV and -12.55 mV, suggesting successful surface CM coating and MSA-2 loading.

The release behavior of MSA-2 from CM/MSA-2@VLPSi was investigated in PBS. The release rates of MSA-2 increased rapidly in the first 8 h and gradually slowed down (Figure S1b and c, Supporting information). After 8 h in PBS, about 86.9% and 91.0% of MSA-2 released from CM/MSA-2@VLPSi-5 and CM/MSA-2@ VLPSi-30. When the time prolonged to 24h, they were about 90.1% and 94.4%, respectively. The results revealed that CM/MSA-2@VLPSi possessed sustained release profile of MSA-2 under a physiological environment, which is essential for activating long-lasting immune response.

### 2.2 Intracellular Ca²^+^ dynamics regulated by VLPSi-activated mechanosensitive Piezo1 channel

Intracellular Ca²^+^ homeostasis is regulated in live cells, where cytosolic Ca²^+^ concentrations are maintained at levels substantially lower than those in the extracellular environment. Multiple intracellular Ca²^+^ reservoirs, particularly the endoplasmic reticulum (ER), cooperate with plasma membrane channels, pumps, and Ca²^+^ -binding proteins to sustain this dynamic equilibrium^33^. To investigate whether VLPSi modulate Ca²^+^ dynamics, intracellular Ca²^+^ levels were monitored using Fluo-4–based fluorescence imaging following stimulation with VLPSi with varying spike lengths (Figure 2a). Compared with the control group, VLPSi-5 increased the proportion of responsive cells from 43.5% to 88.2%. Notably, VLPSi-30 further increased the percentage of responsive cells to 94.1% (Figure 2b). Consistently, both VLPSi-5 and VLPSi-30 induced increased intracellular Ca²^+^ levels compared with the control, and distinct kinetic differences were observed (Figure 2c, S2a and Video S3, Supporting Information). During the early phase (0–300 s), no significant Ca²^+^ increase was detected in the control and VLPSi-5 groups (Figures 2d-f). In contrast, VLPSi-30 triggered a rapid and obviously Ca²^+^ elevation immediately after stimulation, specifically at the 300 s time point, indicating a spike length–dependent mechanosensitive response. In the later phase (300–900 s), intracellular Ca²^+^ levels in the VLPSi-30 group continued to rise at a significantly faster rate than the VLPSi-5 group, suggesting sustained Ca²^+^ influx beyond the initial mechanical trigger. We further selected representative cells to visualize intracellular Ca²^+^ influx at different time points (300, 600, 900, 1200, 1500, and 1800 s), thereby evaluating the effect of spike morphology on cellular calcium dynamics (Figure 2g). At 1700 s, ionomycin treatment was applied as a positive control, leading to a rapid increase in intracellular Ca²^+^ in all groups, confirming probe responsiveness and cell viability. Therefore, sustained calcium elevation following the initial mechanical trigger may involve the activation of secondary calcium signaling cascades downstream. These results of both the proportion of active cells and the fluorescence signal amplitude demonstrate that the VLPSi effectively modulate Ca²^+^ signaling in a spike length–dependent manner, indicating that nanoscale surface topology plays a critical role in regulating cellular Ca²^+^ dynamics.

**Figure 2.**
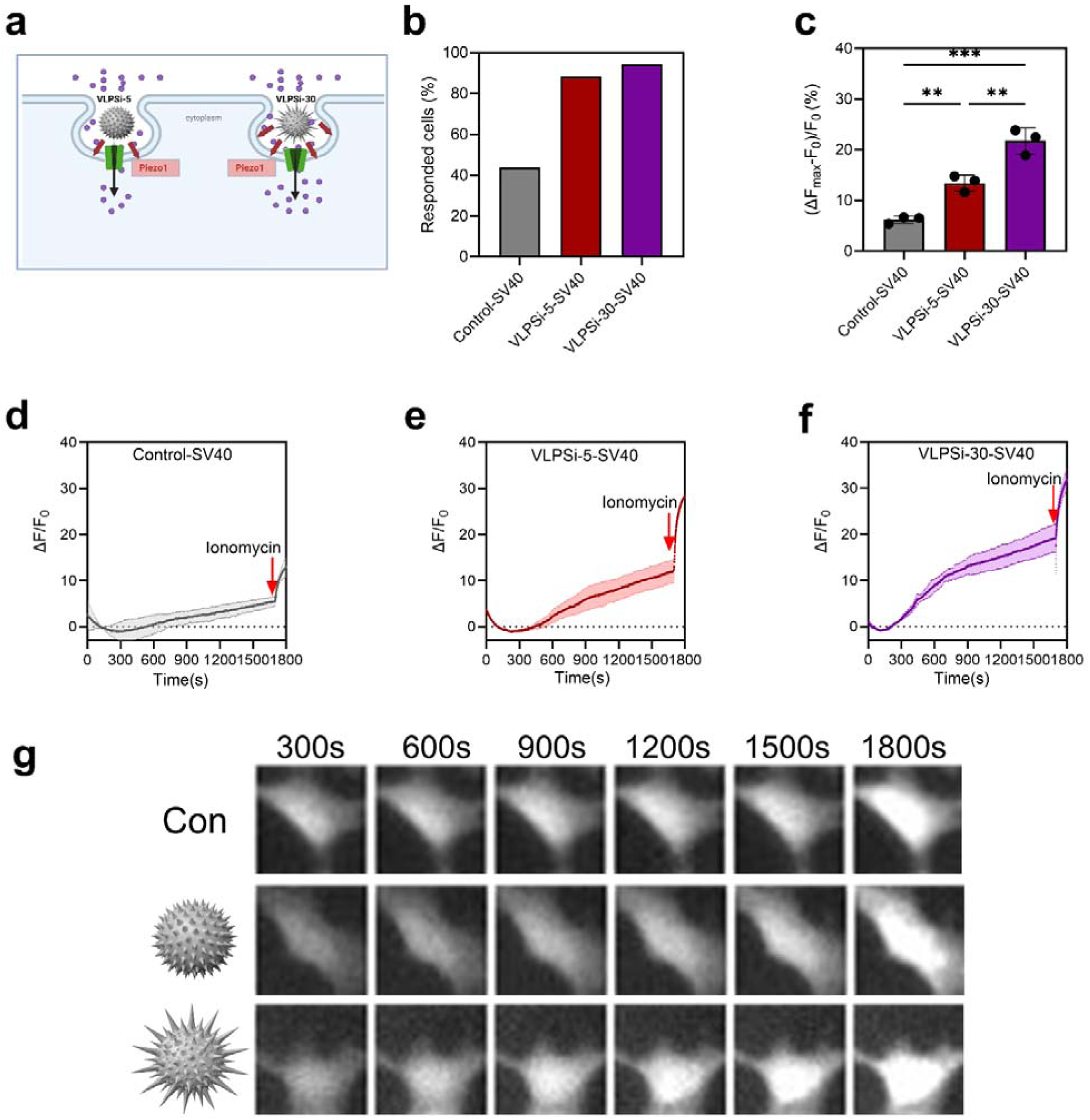
Monitoring of intracellular calcium dynamics based on Fluo-4 fluorescence in SV40 cells. a. The schematic illustration of Ca^2+^ influx by VLPSi-5 and VLPSi-30. b. The proportion of responsive cells caused by VLPSi-5 and VLPSi-30. c. Bar graphs for fluorescent Ca^2+^ signals, the data are presented as mean ± SD; d. Fluorescent Ca^2+^ signals (F-F_o_/F_o_) correspond to control, batches n = 3. e. Fluorescent Ca^2+^ signals (F-F_o_/F_o_) correspond to VLPSi-5. f. Fluorescent Ca^2+^ signals (F-F_o_/F_o_) correspond to VLPSi-30. g. Representative images of control, VLPSi-5 and VLPSi-30 pretreated with SV40 cells at varying times, respectively.

Mechanically activated ion channels, such as Piezo1, are known to function as stiffness-sensitive mechanosensory in macrophages and play critical roles in regulating polarization and inflammatory responses^34,35^. Thus, we hypothesize that this rapid Ca²^+^ influx may be associated with mechanical activation of Piezo1 channel by the rigid spiky nanostructure of VLPSi. Thus, intracellular Ca²^+^ levels were further monitored in human induced microglia-like cells (iMGLs) were differentiated from the wild type (WT) iPSC line GM25256 and the Piezo1 knockout (KO) clone F6 following treatment with VLPSi with different spike lengths. Piezo1 forms a trimeric three-bladed propeller-like architecture. The peripheral blade regions are to sense membrane tension, while the intracellular beam acts as a lever-like mechanical transduction element that couples blade motion to the central pore (Figure 3a). In WT cells, both VLPSi-5 and VLPSi-30 significantly increased intracellular Ca²^+^ influx compared with the untreated control, which agreed with the results with SV40 cells (Figure 3b-c). In contrast, Piezo1 knockout cells substantially decrease the calcium response triggered by VLPSi treatment. Both VLPSi-5 and VLPSi-30 induced only minimal increases in intracellular Ca²^+^ in Piezo1-knock out cells, with no significant differences compared with the control, indicating that Piezo1 is the dominant but not the exclusive contributor to VLPSi-induced calcium signaling (Figure 3d). It demonstrates that VLPSi-induced intracellular calcium influx is dependent on Piezo1-mediated mechan-transduction. In WT cells, VLPSi-treated groups exhibited a gradual and sustained increase in intracellular Ca²^+^ over time, with the strongest response observed in the VLPSi-30 group (Figure 2b, support information). By contrast, Piezo1 KO cells showed markedly weakened calcium dynamics following VLPSi exposure (Figure 2c, support information). The addition of ionomycin at 1700 s, all groups exhibited a rapid increase in intracellular Ca²^+^, indicating that the calcium signaling remained functional and confirming cell viability. Collectively, these results demonstrate that VLPSi induces intracellular calcium influx in a spike length-dependent manner mainly through Piezo1-mediated mechanotransduction, with longer spikes triggering stronger activation.

**Figure 3.**
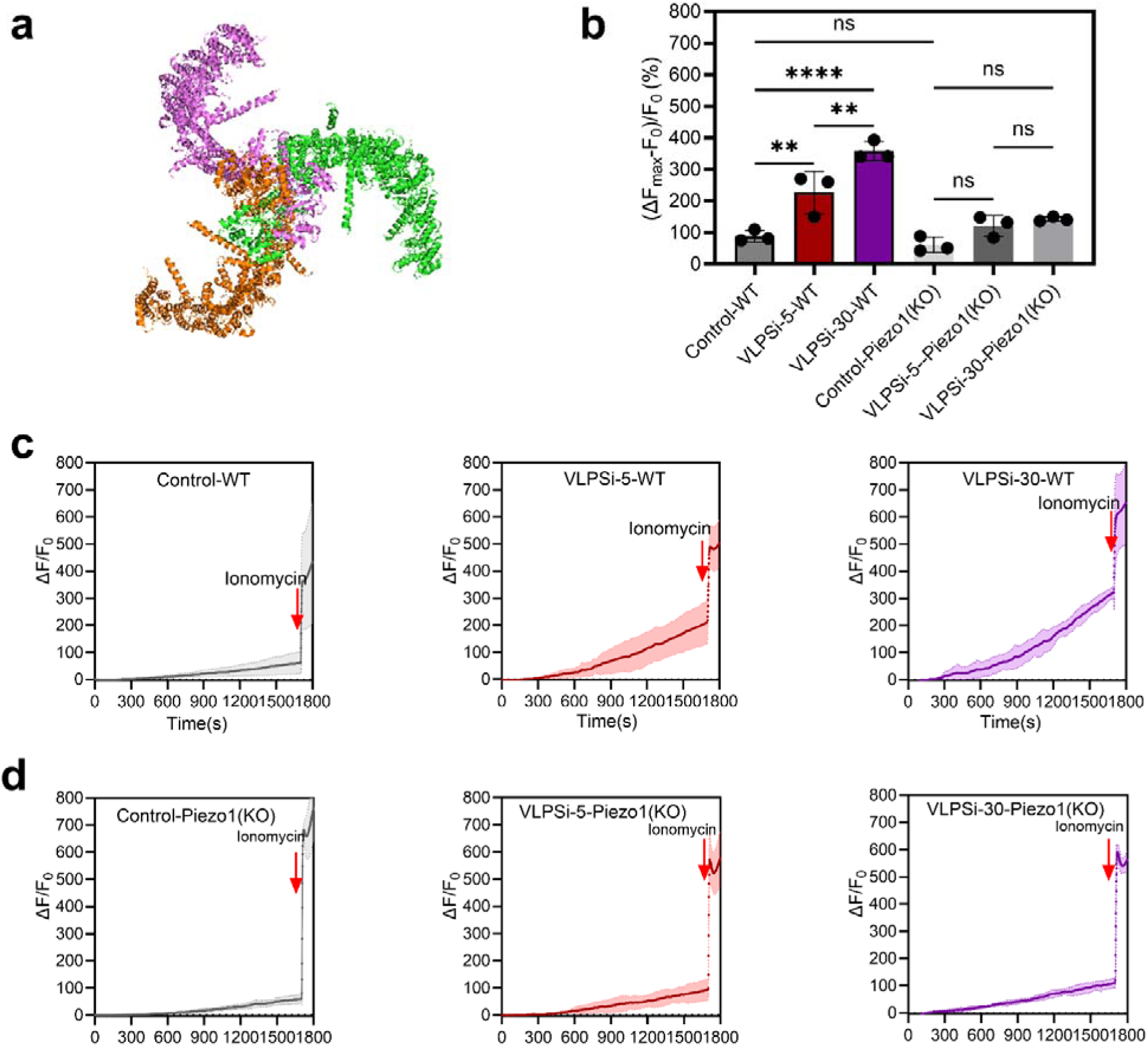
Monitoring intracellular calcium dynamics based on Fluo-4 fluorescence in WT and the Piezo1 knockout (KO) cells. a. The structural model of mouse Piezo1. b. bar graphs for fluorescent Ca^2+^ signals, the data are presented as mean ± SD; c. Fluorescent Ca^2+^ signals (F-F_o_/F_o_) correspond to control, VLPSi-5, VLPSi-30 in WT cells, batches n = 3. d. Fluorescent Ca^2+^ signals (F-F_o_/F_o_) correspond to control, VLPSi-5, VLPSi-30 in Piezo1(KO) cells.

### 2.3 Cellular uptake and cytotoxicity *in vitro*

To evaluate the internalization of VLPSi by RAW264.7 cells, we labeled the NPs with FITC. FITC-labeled NPs displayed green color, while CellMask™-labeled cell membranes showed a distinct red color (Figure 4a). The fluorescence images showed a higher intensity of the FITC signal within RAW264.7 cells that were incubated with FITC-labeled CM/MSA-2@VLPSi-30 compared with those incubated with FITC-labeled CM/MSA-2@VLPSi-5. This suggested that the longer spikes of the VLPSi enhances their cellular internalization.

**Figure 4.**
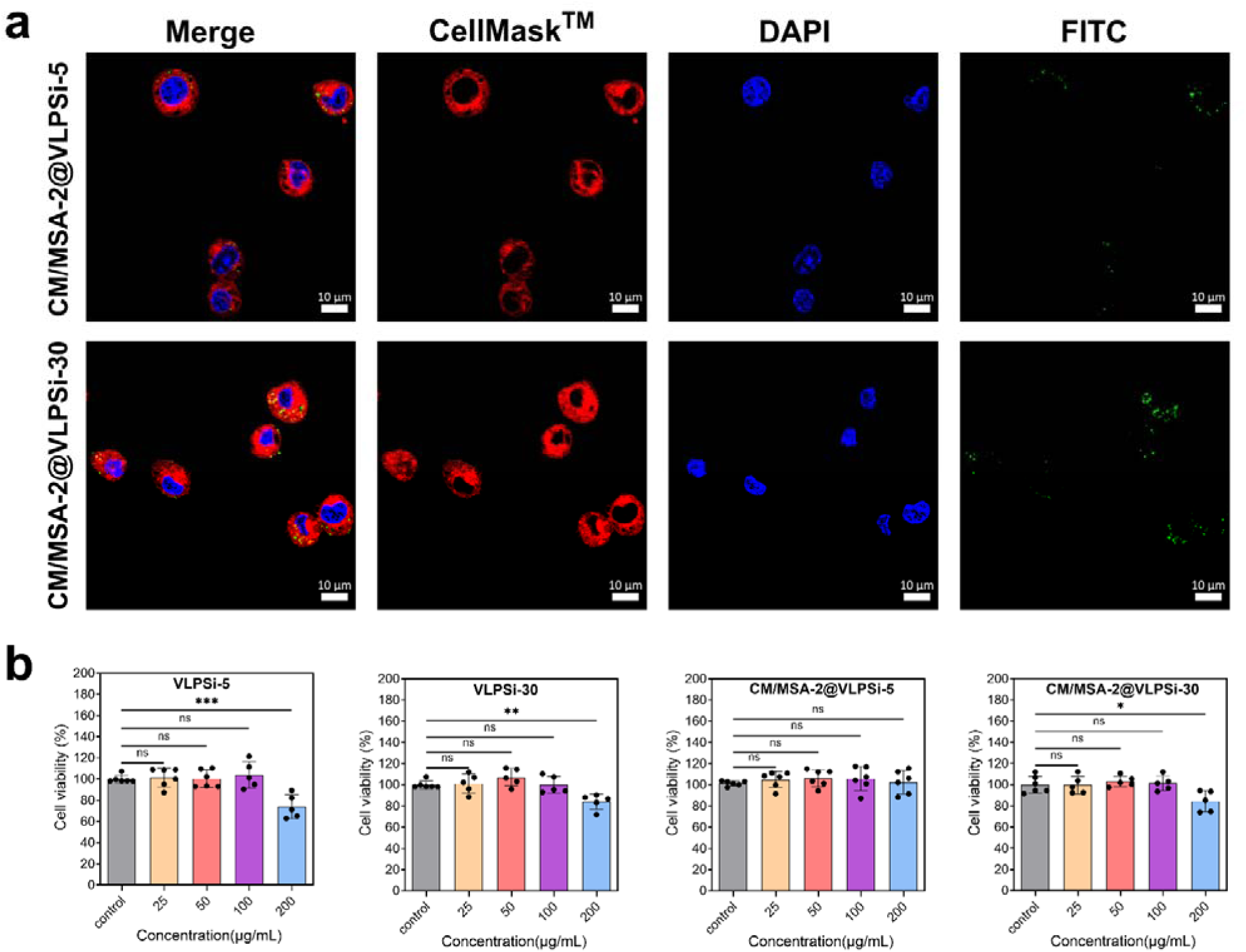
The cytotoxicity and uptake with VLPSi and CM/MSA-2@VLPSi. a. Fluorescence images of Raw264.7 cells treated with FITC-labeled CM/MSA-2@VLPSi of varying surface spike lengths. Nuclei are stained in blue, nano adjuvant in green and cell membrane in red. The scale bar is 10µm.b. Cell viability of VLPSi-5, VLPSi-30, CM/MSA-2@VLPSi-5 and CM/MSA-2@VLPSi-30 treated with different concentrations after 24h. The data represents mean□±□SD. *, ** and *** represent statistically significant differences P < 0.05, P < 0.01 and P < 0.001, respectively.

To evaluate the safety of various VLPSi *in vitro*, the biocompatibility of VLPSi was investigated. The cytotoxicity of the samples VLPSi-5 and VLPSi-30 against RAW264.7 cells was negligible when the concentration was below 200 μg/mL (Figure 4b). A minor toxicity was observed if the concentration increased to 200 μg/mL. Nevertheless, CM/MSA-2@VLPSi-5 did not show any cytotoxicity effect at the same concentration because CM coating and PEGylation enhanced biocompatibility. On the other hand, cell viability was about 84.39% when treated with CM/MSA-2@VLPSi-30 at the same condition due to longer spike, which caused higher cell uptake. The concentration of NPs was lower than 100 μg/mL in the following tests because the NPs did not cause cytotoxicity to cells during experiments.

### 2.4 STING pathway activation

To further explore activation of the STING pathway (Figure 5a), the mouse macrophage cell line RAW 264.7 was treated with CM/MSA-2@VLPSi-5 and CM/MSA-2@VLPSi-30 at a concentration of 100 μg/mL. We further investigated the activation of the STING signaling pathway by western blot analysis. Compared with the control group, the levels of STING, TBK1 and IRF3 were increased in RAW264.7 cells after treated with CM/MSA-2@VLPSi-5 and CM/MSA-2@VLPSi-30 (Figure 5b). These findings indicate that CM/MSA-2@VLPSi can activate the STING pathway and trigger the STING downstream signaling cascade, leading to phosphorylation of TANK-binding kinase 1 (TBK1) and IFN regulatory factor 3 (IRF-3) and subsequently initiating a type I interferon signaling cascade^36^. Western blot assay further verified that CM/MSA-2@VLPSi -30 treatment promoted phosphorylation of NF-κB subunit p65 (p-P65) protein in macrophage cells. The pro-inflammatory functions of M1 macrophages, particularly those mediated by NF-κB signaling, have been extensively investigated^37^. After STING activation, TANK-binding kinase 1 (TBK1) and its homolog IκB kinase ε (IKKε) activate the IKK complex, leading to the subsequent activation of the transcription factor nuclear factor κB (NF-κB). NF-κB cooperates with interferon regulatory factor 3 (IRF3) to drive the robust transcription of type I interferons and other pro-inflammatory cytokines, thereby amplifying innate immune responses^38^. IFN-β cytokines not only play an important role in directly antitumor effects but also facilitate DC maturation and bridge innate and adaptive immunity^39^. To evaluate this function, RAW264.7 were exposed to either CM/MSA-2@VLPSi-5 and CM/MSA-2@VLPSi-30. Both can significantly up-regulated major expression of IFN-β because of MSA-2 loading, whereas CM/MSA-2@VLPSi -5 had only a modest effect compared to CM/MSA-2@VLPSi -30 (Figure 5c). Thus, the spike length of VLPSi can also affect the IFN-β secretion of cytokines.

**Figure 5.**
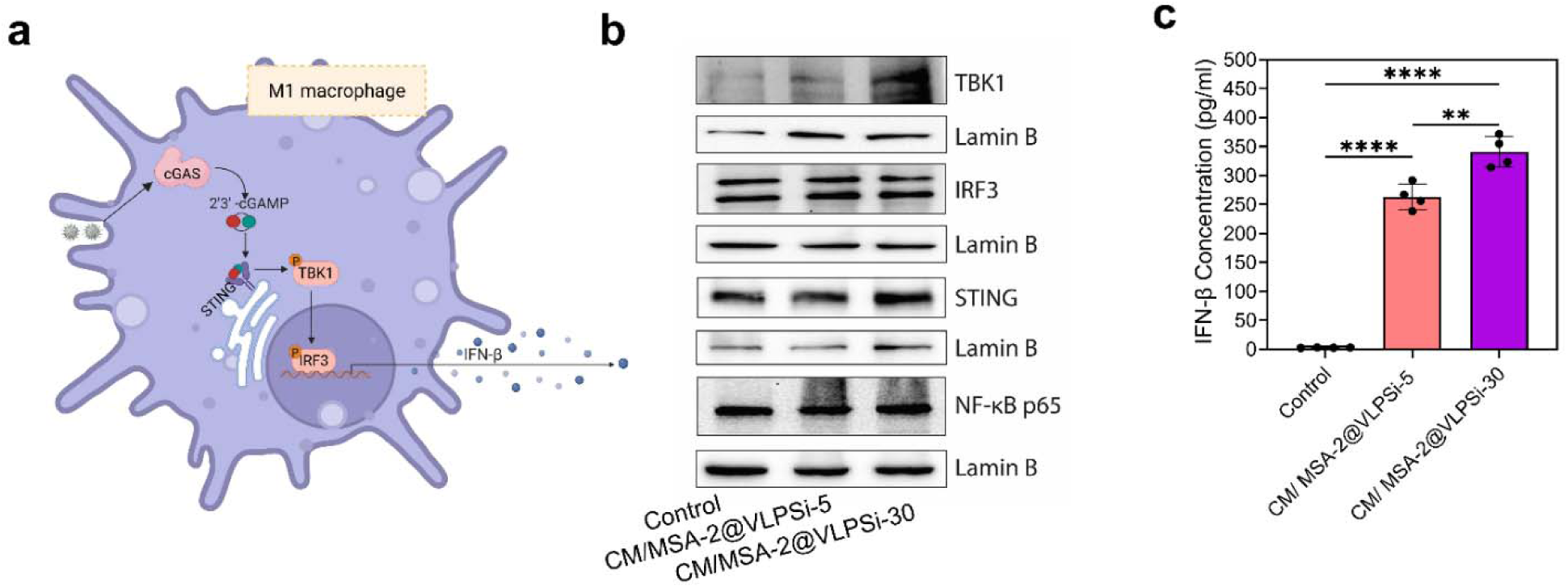
Anti-inflammatory effect of CM/MSA-2@VLPSi-5 and CM/MSA-2@VLPSi-30. a. The schematic illustration of molecular mechanisms involved in M1 macrophage polarization. b. Flow cytometry quantification of CD86-positive with 100ug/mL CM/ MSA-2@VLPSi-5, 30. c Flow cytometry quantification of CD206-positive cells. d. Western blot analysis of proteins related to STING signaling pathway in RAW264.7 cells. e. Levels of IFN-β in culture media of RAW264.7 cells with 100ug/mL CM/MSA-2@VLPSi-5 and CM/MSA-2@VLPSi-30 treatment for 24 h were determined by using the ELISA kit. The data represents mean□±□SD, * P < 0.05; ** P < 0.01; *** P < 0.001.

### 2.5 Macrophage polarization

Macrophages are innate immune cells characterized by their functional characteristics, known as M0 (resting), M1 (pro-inflammatory), or M2 (anti-inflammatory), while Lipopolysaccharide (LPS) and IFN-γ are an inducer of the M1 macrophage phenotype^40^. Accordingly, we used IFN-γ 20 ng/mL and LPS 100 ng/mL as positive control to polarize RAW 264.7 cells for M1 macrophage phenotype. CM/MSA-2@VLPSi-5 and CM/MSA-2@VLPSi -30 were incubated with cells and quantify their polarization efficacy via flow cytometry analysis and RT-qPCR (Figure 6a).

**Figure 6.**
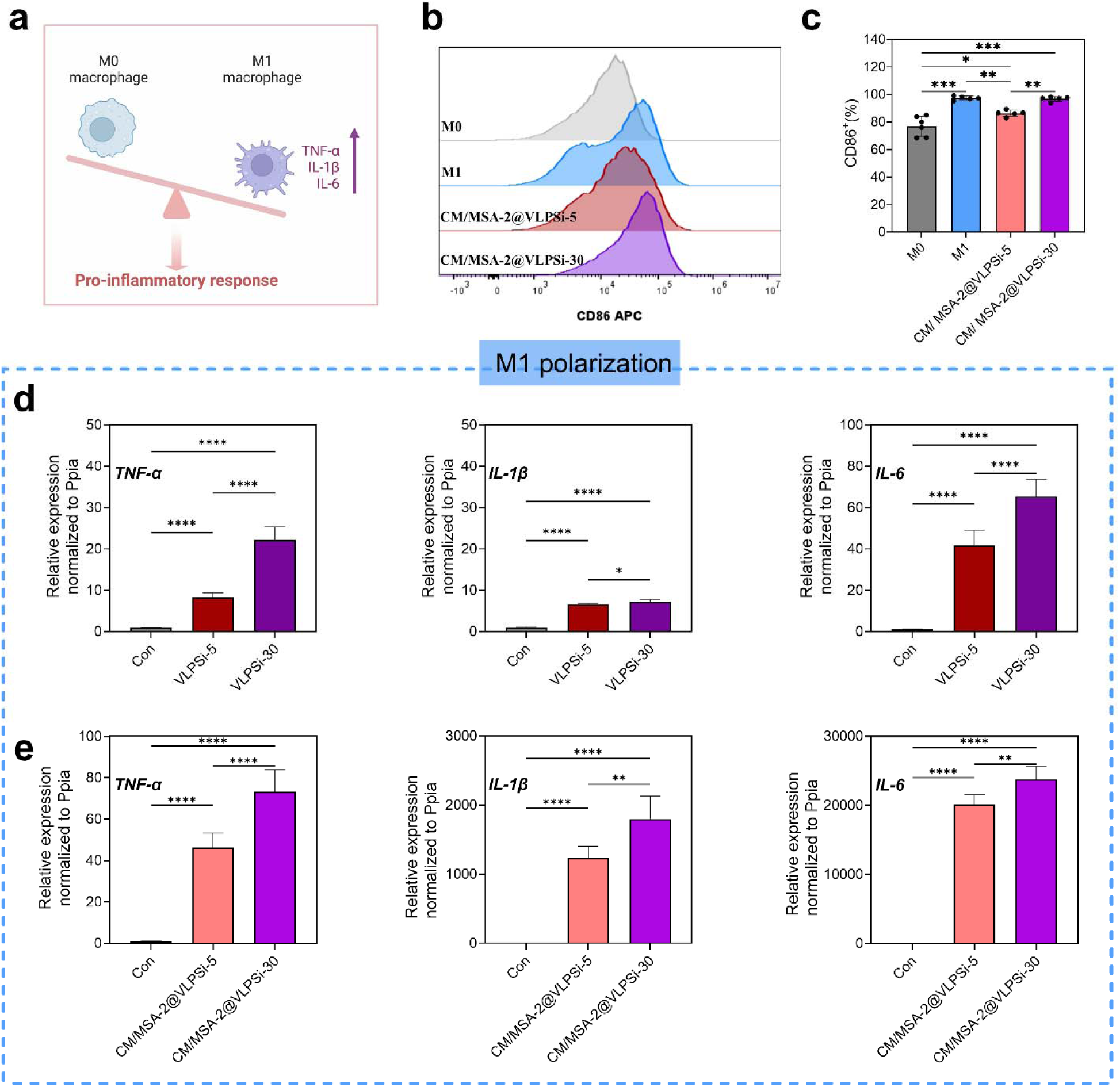
Influence of CM/ MSA-2@VLPSi-5 and CM/ MSA-2@VLPSi-30 on the regulation of macrophage polarization. a. The schematic illustration of macrophage polarization. b. Flow cytometry quantification of CD86-positive with 100ug/mL CM/ MSA-2@VLPSi-5, 30. c. Representative histograms of CD86-positive with particles. M0 represents control group, M1 represents the cells that were treated with IFN-γ 20 ng/mL and LPS 100 ng/mL. d. Expression level of M1 macrophage markers TNF-α, IL-1β and IL-6 in VLPSi-5 and VLPSi-30. e. Expression level of M1 macrophage markers TNF-α, IL-1β and IL-6 in CM/MSA-2@VLPSi-5 and CM/MSA-2@VLPSi-30. The data represents mean□±□SD, * P < 0.05; ** P < 0.01; *** P < 0.001.

The proportion of CD86-positive macrophages was significantly increased by the addition of positive control LPS+ IFN-γ. Notably, the CM/MSA-2@VLPSi-30 group exhibited a higher proportion of CD86-positive macrophages than the CM/MSA-2@VLPSi-5 group (Figure 6b). Quantification of the percentage of CD86^+^ cells revealed that CM/MSA-2@VLPSi-5 and CM/MSA-2@VLPSi-30 exhibited pro-inflammatory potencies, with the M1 macrophage percentage being approximately 86.3% and 96.8%, respectively (Figure 6c). Quantification of the percentage of CD206^+^ cells revealed that there was no significant difference between CM/MSA-2@VLPSi-5 and CM/MSA-2@VLPSi-30 (S4a-b, Supporting information). These results indicated that CM/MSA-2@VLPSi-30, especially the longer spike length, could effectively promote the pro-inflammatory M1-type macrophages.

Therefore, the effects of VLPSi on the expression levels of M1 and M2 key factors were further investigated by using RT-qPCR. M1-type macrophage-associated pro-inflammatory levels, such as TNF-α, IL-1β, and IL-6, were increased after stimulation. As expected, VLPSi-30 and CM/MSA-2@VLPSi-30 more significantly increased the levels of these factors than VLPSi-5 and CM/MSA-2@VLPSi-5 (Figure 5d-e). Activation of the NF-κB pathway was accompanied by increased expression of pro-inflammatory cytokines, including TNF-α, IL-1β, and IL-6, together with macrophage activation toward a pro-inflammatory phenotype^41^. Additionally, the expression levels associated with M2-type macrophages were also examined, such as Arg-1, IL-4 and TGF-β (S4c-d, Supporting information). The addition of NP didn’t change IL-4, Arg-1 and TGF-β expression level among VLPSi-5, VLPSi-30, CM/MSA-2@VLPSi-5, CM/MSA-2@VLPSi-30 groups, indicating there was no polarization towards the M2 phenotype. Taken together, these results indicate that the longer spike length (30nm) promotes the higher expression of proinflammatory biomarkers and have the effect on the pro-inflammatory M1-type macrophages.

## Conclusions

In summary, we demonstrated that virus-like nanotopography plays an active role in polarizing M1 macrophages and subsequently developed a biomimetic cancer cell membrane-coated, STING agonist-loaded virus-like mesoporous silica nanoplatform (CM/MSA-2@VLPSi) for macrophage programming. Compared to the VLPSi with 5 nm spikes (VLPSi-5), VLPSi-30 with longer spike exhibited significantly enhanced cellular internalization. Mechanistically, the rigid spiky topography facilitates the opening of mechanosensitive Piezo1 channels and disrupted intracellular Ca²^+^ homeostasis, resulting in increased Ca²^+^ influx when the cells were treated with VLPSi-30. Beyond Ca²^+^ -mediated immunomodulation, CM/MSA-2@VLPSi directly activated the STING activation through MSA-2 delivery. The expression levels of STING, TBK1, IRF3, NF-κB, and IFN-β were spike length– dependent, with significantly enhanced activation when the VLPSi with longer spike. Piezo1-mediated Ca²^+^ influx contribute to NF-κB activation, whereas MSA-2 activates the STING pathway. The convergence of Ca²^+^ -dependent NF-κB signaling and STING-mediated inflammatory signaling may synergistically promote macrophage M1 polarization. Coating with IFN-γ treated CM thereby enhanced biocompatibility of the nanoparticles and antigen presentation for activation of immune cells. Consequently, the VLPSi with longer spikes markedly upregulated the expression of pro-inflammatory cytokines, including TNF-α, IL-1β, and IL-6, thereby promoting M1 macrophage polarization. These findings reveal that nanoscale physical cues can function as upstream regulators of innate immune signaling and suggest that the coordinated integration of mechanical stimulation, inflammatory activation, and antigen presentation represents a promising strategy for enhancing antitumor immunity. More broadly, this work provides new insight into the interplay between mechanotransduction and immune signaling and establishes a foundation for the rational design of next-generation biomimetic immunotherapies.

## Supporting information

Activation of Piezo1 by Virus-like Nanospikes-S3-control

Activation of Piezo1 by Virus-like Nanospikes-S3-VLPSi-5

Activation of Piezo1 by Virus-like Nanospikes-S3-VLPSi-30

## Acknowledgments

This work was supported by Research Council of Finland (Grant no. 356056), program of China Scholarship Council, and Cancer Foundation Finland (Grant No. 230130). SIB Labs, UEF Cell and the Tissue Imaging Unit were acknowledged for technical support. We also acknowledge Medicinal Chemistry and Organic Synthesis Facilities (MCOS) of the Drug Discovery, Chemical Biology (DDCB) platform, as part of the Biocenter Kuopio and Biocenter Finland, as well as laboratory technician Tiina Koivunen for assistance with the sample preparation.

