## Supplementary Information for "Mechanical Activation of Piezo1 by Virus-like Nanospikes to Potentiate STING-driven Macrophage Reprogramming"

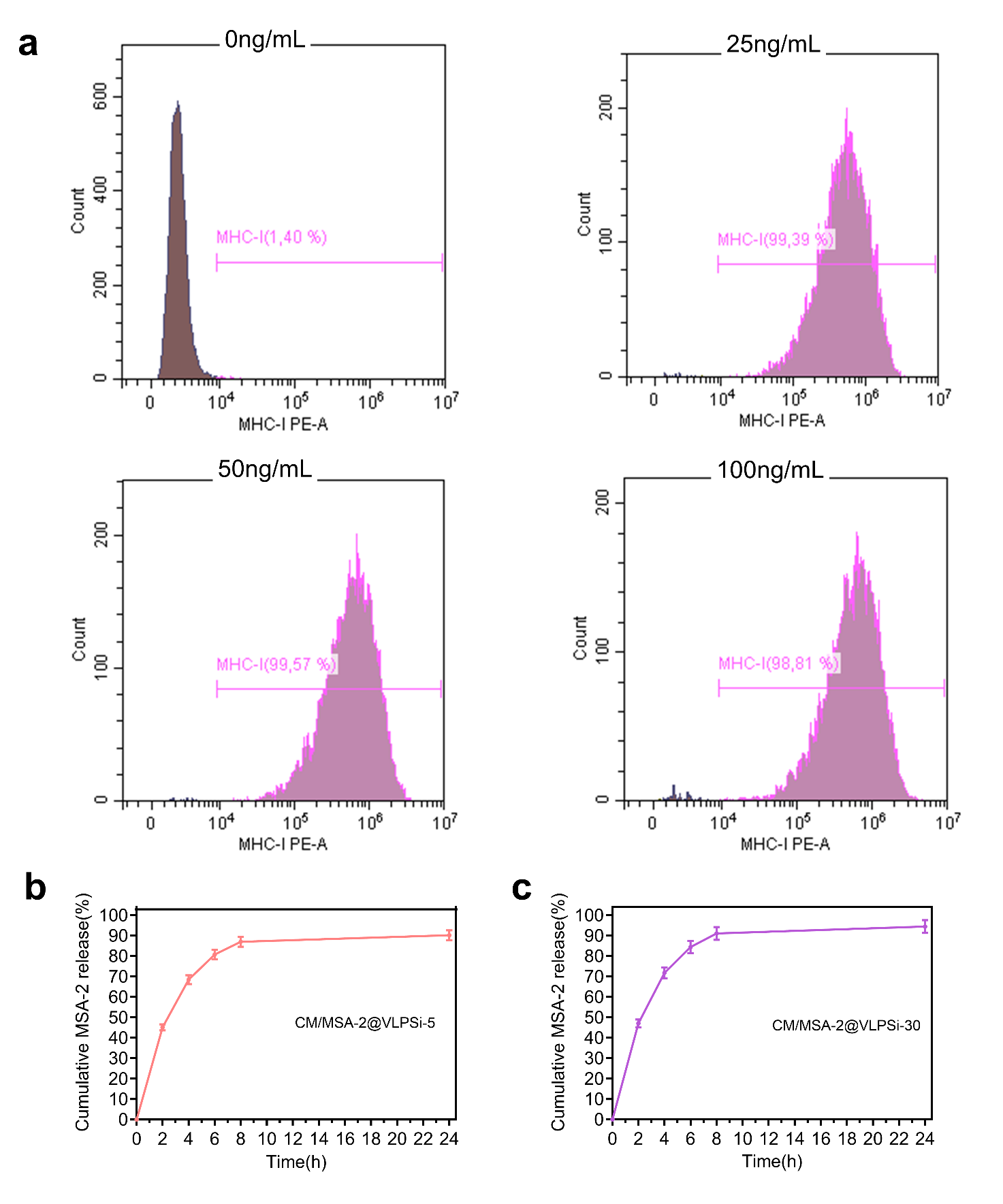

S1. a. Flow cytometry of three different concentrations of IFN-γ to stimulate MHC І expression on the B16-OVA cell surface. b. The release of. MSA-2 from CM/MSA-2@VLPSi-5 in PBS at 0, 2, 4, 6, 8, 24h. c. The release of. MSA-2 from CM/MSA-2@VLPSi-30.

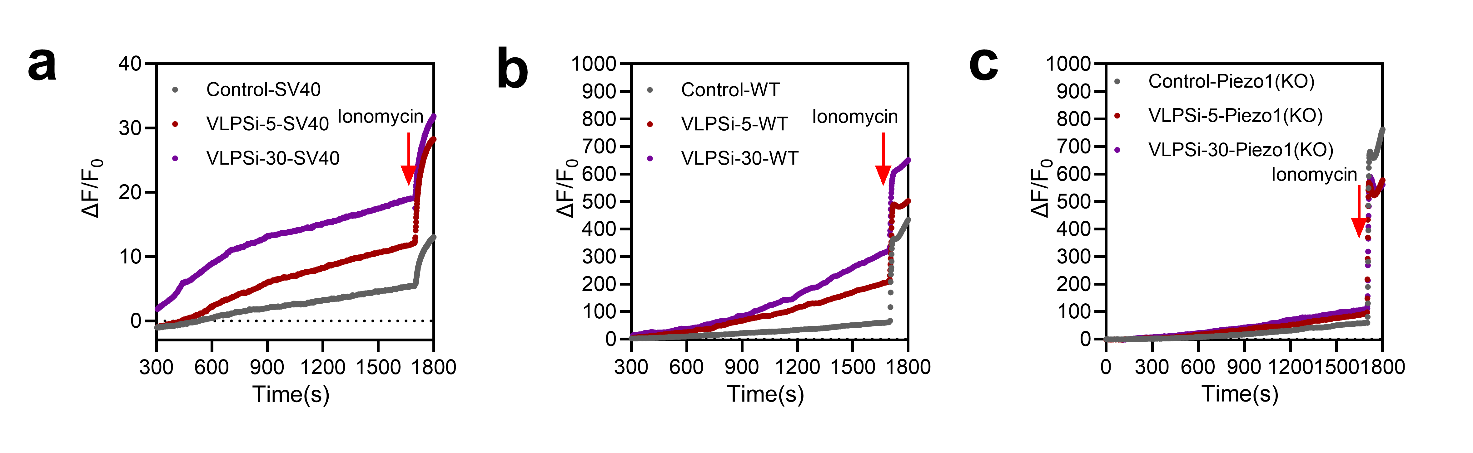

S2. a. Fluorescent Ca2+ signals (F-Fo/Fo) during mechanical stimulation in SV40 cells with control, VLPSi-5 and VLPSi-30, batches n = 3. b. Fluorescent Ca2+ signals (F-Fo/Fo) in WT cells. c. Fluorescent Ca2+ signals (F-Fo/Fo) in Piezo1(KO) cells. d. The dynamic of spatiotemporal distribution of intracellular Ca2+ in adherent cells incubated with VLPSi-5 and VLPSi-30. The recordings extended over 30 minutes, with image acquisition at 1-second intervals. The playback frame rate is 200 frames per second.

Supporting Videos

S3. The dynamic of spatiotemporal distribution of intracellular Ca2+ in adherent cells incubated with control, VLPSi-5 and VLPSi-30. The recordings extended over 30 minutes, with image acquisition at 1-second intervals. The playback frame rate is 200 frames per second.

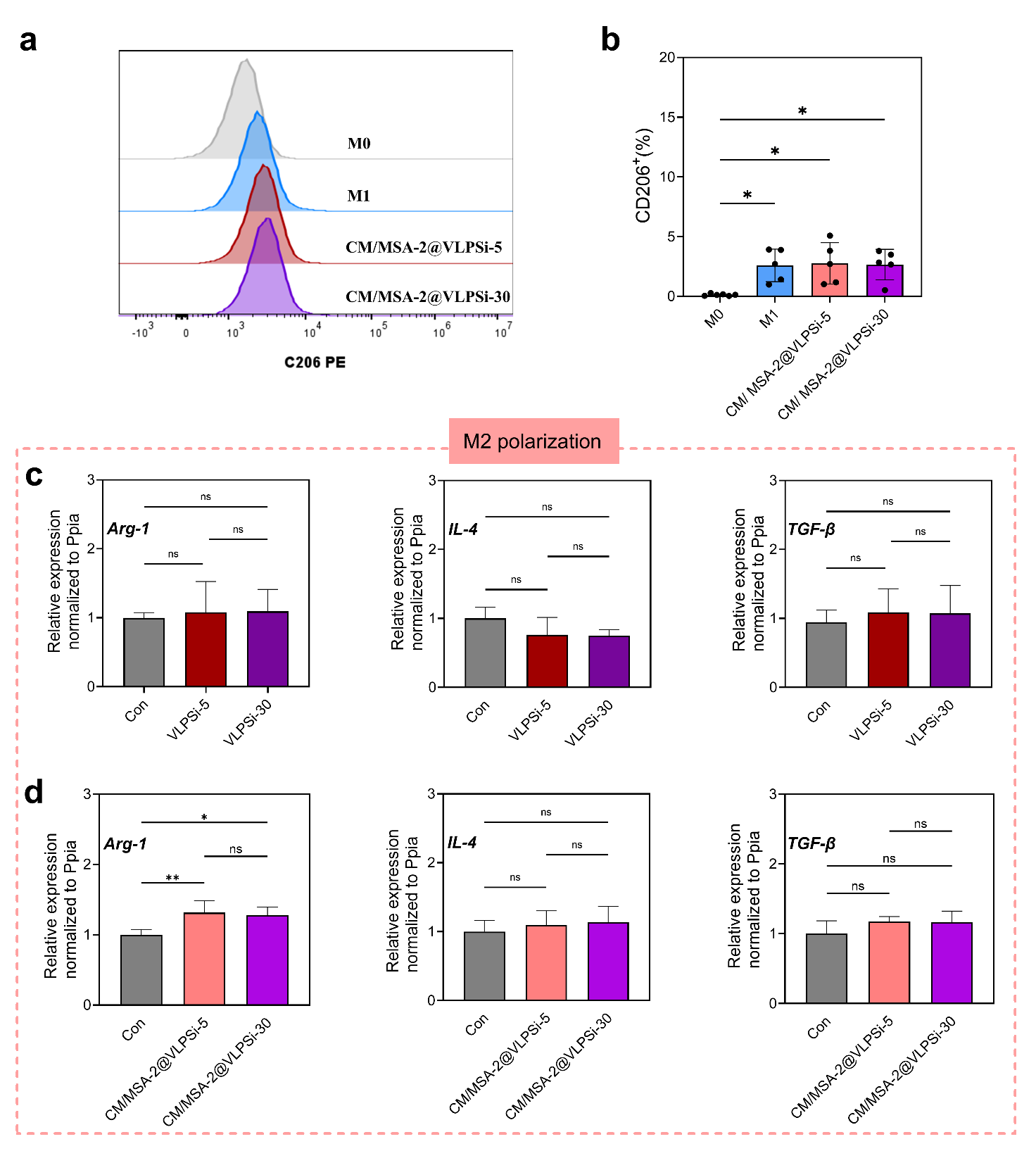

S4. a. Flow cytometry quantification of CD206-positive with 100ug/mL CM/ MSA-2@VLPSi-5, 30. b. Representative histograms of CD206-positive with particles. M0 represents control group, M1 represents the cells that were treated with IFN-γ 20 ng/mL and LPS 100 ng/mL. c. Expression level of M2 macrophage markers Arg-1, IL-4 and TGF-β. d. Expression level of M2 macrophage markers Arg-1, IL-4 and TGF-β. The data represents mean ± SD, * P < 0.05; ** P < 0.01; *** P < 0.001.

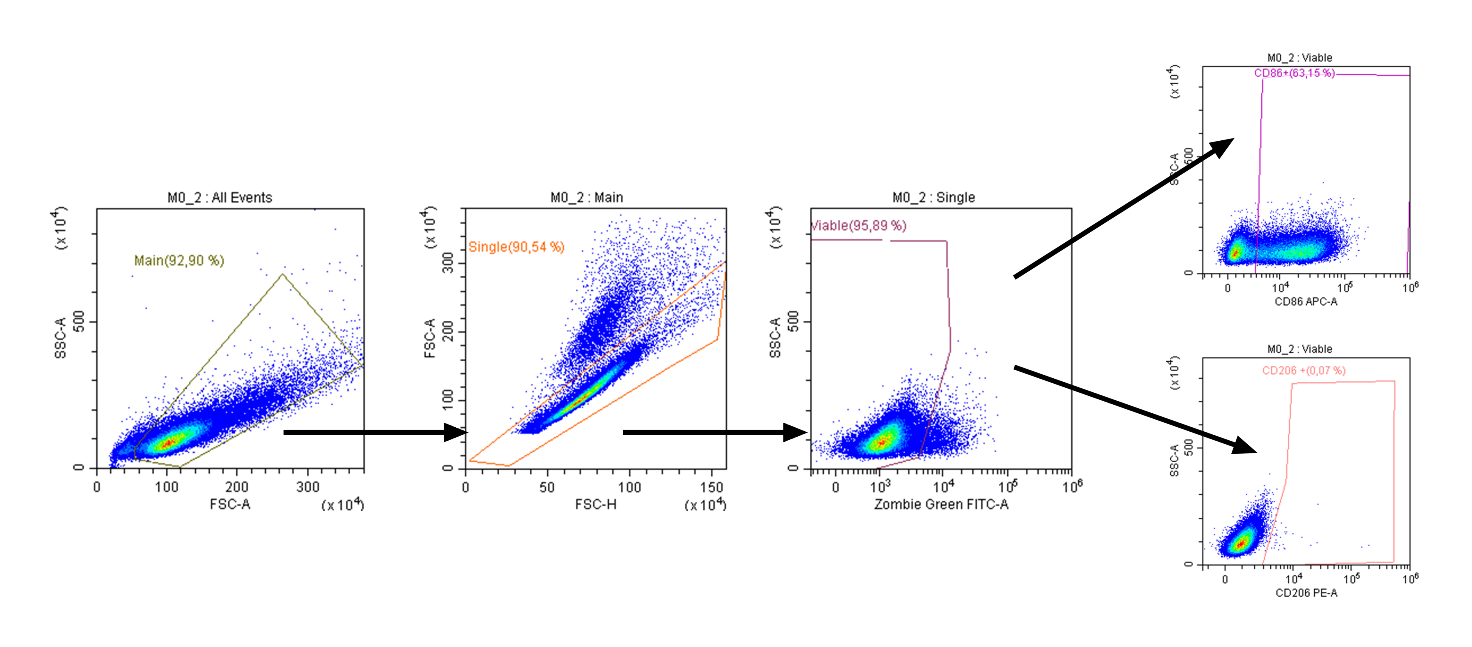

S5. Representative example for the gating strategy in RAW264.7 cells.
